# Kinetics of cortisol and cortisone binding to corticosteroid binding globulin and albumin *in vivo*

**DOI:** 10.64898/2026.04.14.718600

**Authors:** Aurora K Authement, Abhinav Nath, Katya B Rubinow, John K Amory, Nina Isoherranen

## Abstract

Cortisol is a major endogenous glucocorticoid that regulates numerous physiological processes. In plasma, cortisol and its inactive metabolite cortisone bind to corticosteroid-binding globulin (CBG) and albumin, leaving only the unbound fraction available for receptor activation and metabolism. Changes in ligand or protein concentrations alter unbound fractions. Existing binding equations are difficult to extend to multi-ligand, multi-protein systems and do not readily capture competitive endogenous binding interactions. The goal of this study was to develop a plasma protein binding model that quantitatively describes binding species and predicts unbound concentrations across physiological states.

Total and unbound cortisol and cortisone, CBG and albumin were measured in plasma from healthy premenopausal women (n=13) at baseline and after 7 days of 30 mg hydrocortisone treatment. Reversible 1:1 binding models were implemented in COPASI and MATLAB/Simulink, and dissociation constants (K_d_) were estimated by fitting binding models to observed unbound concentrations. A model describing simultaneous binding of cortisol and cortisone to CBG and albumin yielded *in vivo* K_d_ values for cortisol:CBG, cortisone:CBG, cortisol:albumin, and cortisone:albumin of 0.0130 µM, 0.169 µM, 172 µM, and 519 µM, respectively. Model predictions agreed with observed unbound cortisol and cortisone, and bootstrap resampling confirmed stable K_d_ estimates.

This work provides a quantitative framework for predicting unbound cortisol and cortisone across physiological and disease states by accounting for both changes in ligand and protein concentrations. This enables extrapolation without reparameterization and supports exploration of conditions such as pregnancy, adrenal insufficiency, and liver disease, informing interpretation of altered cortisol concentrations in these populations.

**Significance statement:** This work establishes a framework to predict *in vivo* cortisol and cortisone binding. The developed model was applied to predict unbound cortisol and cortisone concentrations in physiological and pathophysiological states and can be integrated into pharmacokinetic models. Our analysis demonstrates that cortisol and cortisone binding affinities estimated in the native plasma environment differ from those measured using purified proteins. These differences have important implications for predicting and analyzing unbound cortisol concentrations.

**Graphical Abstract:** 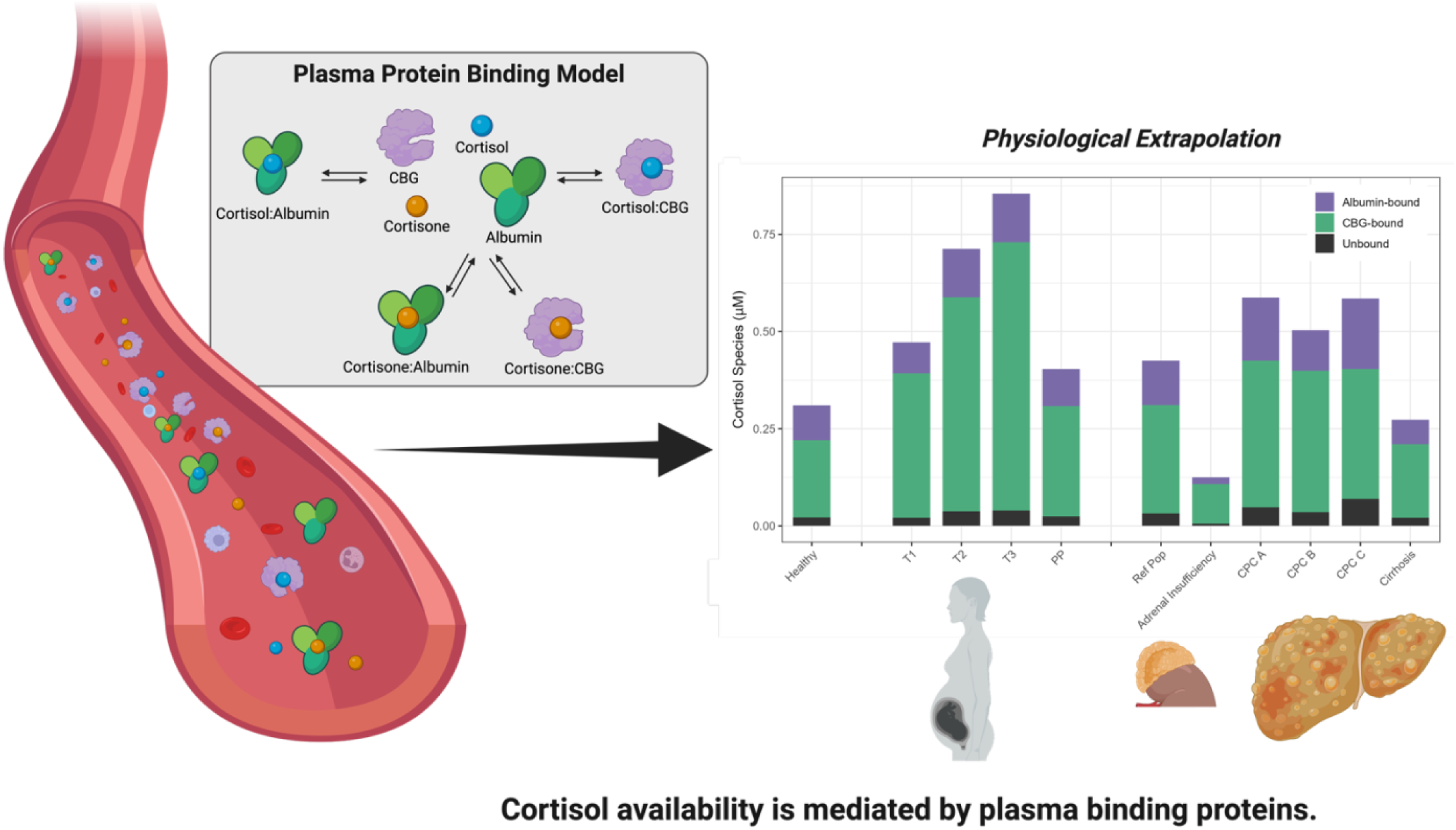

Created in BioRender. Authement, A. (2026) https://BioRender.com/zl1bg0k

## 1. Introduction

Cortisol is the most potent endogenous glucocorticoid with critical roles in metabolism, immune activity, and the physiological stress response (1). Although total serum cortisol concentration is often measured clinically, only the unbound fraction is biologically active and diffuses across cell membranes to interact with the glucocorticoid receptor (2,3). In circulation, cortisol is predominantly bound to corticosteroid binding globulin (CBG) and, to a lesser extent, albumin (4,5). CBG provides high-affinity, low-capacity binding, while albumin contributes low-affinity, high-capacity interactions. The low capacity, high affinity binding of cortisol to CBG causes the unbound fraction (f_u_) of cortisol to be concentration-dependent (6). Cortisone, the inactive 11β-hydroxysteroid dehydrogenase 2 (11β-HSD2) metabolite of cortisol (7), also binds to CBG and albumin and thus, cortisone binding may affect cortisol binding kinetics (8,9). The free concentrations of cortisol and cortisone are, therefore, defined by the relative concentrations of binding proteins and endogenous ligands binding to CBG and albumin.

The equilibrium between bound and unbound cortisol is influenced by physiological and pathological changes in binding protein concentrations. Across different physiological and disease states such as pregnancy (10,11), adrenal insufficiency (12–14), and liver disease (15), CBG, albumin, total cortisol and total cortisone concentrations change. As a result, cortisol and cortisone are redistributed among unbound, CBG-bound, and albumin-bound species, altering the fractional concentrations and the balance between bound and unbound cortisol. These perturbations are often described qualitatively in clinical and research studies. Their quantitative impact on free cortisol availability remains poorly defined in mechanistic research assessing glucocorticoid signaling. In research settings, this creates a barrier to interpreting cortisol dynamics mechanistically, especially when investigating hormone-protein interactions, feedback regulation, circadian rhythm, or stress response. Kinetic, quantitative understanding of binding equilibria could enable more precise evaluation of cortisol action across physiologic and pathophysiologic conditions, support comparative studies across populations, and ultimately contribute to establishing context-dependent reference ranges for total serum cortisol. Such ranges would reflect not only rates of cortisol secretion and clearance, but also the plasma binding environment, providing a more biologically informed basis for interpreting cortisol exposure across conditions characterized by altered CBG or albumin.

Established analytical models like the Coolens method (16) or the Dorin-Qualls equation (17), which are respectively based on quadratic and cubic formulas, are extremely challenging or impossible to extend to more complicated systems of interactions. To overcome these limitations, we constructed a plasma protein binding model (PPBM) that accounts for binding of both cortisol and cortisone to CBG and albumin. By numerically modeling the relevant binding equilibria with a system of differential equations, we were able to fit a model to comprehensive dataset of free and total cortisol and cortisone concentrations and CBG and albumin concentrations in 13 healthy volunteers and estimate the *in vivo* binding affinities of cortisol and cortisone to CBG and albumin. The PPBM enables systematic evaluation of how variation in CBG, albumin, cortisol, or cortisone concentrations influences free cortisol availability, as well as prediction of how physiological or pathophysiological changes could impact free cortisol levels.

## 2. Materials and Methods

2.1. *Chemicals and reagents*

Optima LC/MS grade acetonitrile, water, and formic acid were purchased from Thermo Fischer Scientific (Waltham, MA, USA). Charcoal treated blank human serum (DC Mass Spect Gold MSG 4000) was from Golden West Biologics (Temecula, CA, USA). Cortisol, cortisone, and the internal standard 9,11,12,12-d_4_-cortisol (cortisol-d_4_) were purchased from Millipore Sigma (St. Louis, MO, USA).

### 2.2. Clinical study design for the plasma measurement of cortisol, cortisone, CBG, and albumin

Cortisol, cortisone, corticosteroid binding globulin (CBG) and albumin concentrations were measured in plasma samples collected from healthy premenopausal women (n = 13) at baseline conditions and following 7 days of oral hydrocortisone (CORTEF® tablets, Pfizer, New York NY) self-administration (20 mg in the morning and 10 mg in the afternoon). The sample collection was part of a larger crossover clinical study evaluating the impact of endogenous hormones on dronabinol metabolism. The study was approved by the Institutional Review Board at the University of Washington (STUDY0008064), registered at ClinicalTrials.gov (ID NCT04374773) and conducted in accordance with the Declaration of Helsinki principles. None of the participants were taking hormonal contraceptives. Participant demographics and other details regarding inclusion and exclusion criteria can be found at ClinicalTrials.gov. All participants provided written informed consent prior to any study procedures. The control study day reflected endogenous cortisol exposure and diurnal variation in cortisol and cortisone concentrations. The treated day reflected significantly elevated cortisol exposures.

On each study day, serial blood samples were collected at 0, 0.5, 1, 1.5, 2, 3, 4, 6, 8, 10, 12, and 24 hours after the start of the study (geometric mean start time 7:52 AM for control and 7:40 AM for treated). This resulted in 12 samples per participant and a total of 312 plasma samples analyzed. Exact blood draw times are provided in the Supplemental Table 1.

### 2.3. Quantification of cortisol and cortisone

Cortisol and cortisone were analyzed using a previously described method with minor modifications (18). Calibration curves (8 non-zero points) were prepared in Mass Spect Gold charcoal stripped serum at concentrations 0.025-0.900 μM cortisol and 0.015-0.200 μM cortisone with quality control (QC) samples prepared in the same matrix. The QC concentrations are listed in Supplemental Table 2.

To precipitate proteins, 90 µL of ice-cold acetonitrile containing cortisol-d_4_ (0.400 μM) was added into 60 µL of plasma, or calibration or QC standards in a 96-well plate. The plates were centrifuged at 3,000 x g at 4°C for 45 minutes. Supernatants were transferred to clean wells and centrifugation was repeated at 3,000 x g at 4°C for 30 minutes. Supernatant was transferred to a new 96-well plate, and analytes were quantified using Liquid chromatography tandem mass spectrometry (LC-MS/MS) analysis.

Plasma samples were quantified using a Sciex 6500 QTRAP mass spectrometer (Framingham, MA, USA) coupled to an Agilent 1290 Infinity II liquid chromatograph (Santa Clara, CA, USA) operated in negative ion electrospray ionization mode. The mass spectrometer parameters were dwell time 50 milliseconds, curtain gas 35, CAD -2, ionization spray -4500 V, source temperature 500°C, GS1/2 80/60, and EP was -10. Analyte specific parameters DP, CE, and CXP were -30, - 26, and -34 for cortisol and -45, -22, and -15 for cortisone, respectively. The following MRM transitions in the negative ion mode were monitored [M+35Cl−]: *m/z* 407 > 331 for cortisol, *m/z* 405 > 329 for cortisone, and *m/z* 411 > 335 for cortisol-d_4_.

The analytes were separated using a Thermo Hypersil Gold C18 column (100 × 2.1 mm, 1.9 µm, Waltham, MA, USA). The mobile phase flow was 0.5 mL/min, injection volume was 3 µL, and the column temperature was set to 25°C and autosampler was set to 4°C. The mobile phase consisted of (A) water with 0.1% formic acid and (B) acetonitrile with 0.1% formic acid. A gradient elution was used starting from 10% B for 0.5 min, increased to 90% B over 3 min, held for 1.5 min, followed by a return to initial conditions and held for 2 min.

LC-MS/MS data were analyzed using MultiQuant 3.0.3. Analyte concentrations were calculated using peak area ratios relative to cortisol-d_4_. Calibration curves for each analyte were fitted by linear regression and weighted by 1/x. The method was validated according to FDA Bioanalytical Method Validation Guidance (19). All calibration curves and QC samples met acceptance criteria and are reported in Supplemental Table 2.

### 2.4. Quantification of corticosteroid binding globulin and albumin

Plasma corticosteroid binding globulin (CBG) was quantified using a sandwich ELISA (Biovendor, Ashville, NC) according to the manufacturer’s instructions. Each plate included low- and high-quality control samples provided by the manufacturer. Albumin concentrations were measured at enrollment by the clinical laboratory of the University of Washington (UW) Medical Center.

### 2.5. Determination of plasma fraction unbound of cortisol and cortisone

The plasma unbound fractions (f_u_) of cortisol and cortisone were determined by rapid equilibrium dialysis (RED) according to the manufacturer’s recommendations. Plasma (100 µL) was aliquoted into the matrix chamber of the RED Device (Thermo Scientific, Waltham, MA, USA) and 350 µL of 100 mM potassium phosphate (KPi) buffer was added to the buffer chamber. Equilibrium for all analytes was confirmed at 12 hours at 37 °C based on time-course data. At 12 hours, 50 µL of sample from each chamber were collected for analysis. The samples, calibration curve, and quality controls were analyzed by LC-MS/MS as described above. The plasma f_u_ was calculated as the ratio of the unbound concentration of analyte in the buffer chamber to the total concentration in the matrix chamber. The plasma chamber was analyzed as previously described. The buffer chamber quantification was performed in 100 mM KPi at pH 7.4 using eight non-zero points. The concentration range for both cortisol and cortisone was 0.001–0.050 μM, with LQC, MQC, and HQC prepared at 0.003, 0.025, and 0.035 μM, respectively. The maximum f_u_ in each participant was defined as the highest observed f_u_ across the samples. The average f_u_ was calculated as the area under the f_u_–time curve divided by the sampling time (12 hours).

### 2.6. Statistical analysis

Differences in values between the study days were evaluated by paired t-test on log-transformed data using R statistical software (version 4.2; R Core Team 2022). A p-value < 0.05 after false discovery rate (FDR) correction for multiple comparisons was considered significant. Log-normality of the data was assessed by Shapiro-Wilk test. Outliers were identified using Grubbs’ test on the full dataset stratified by analyte and were excluded from further statistical analysis and tabulated data. Five outliers were identified, four in the cortisol dataset and one in the cortisone dataset. If shown in figures, outliers are denoted with an “X” symbol.

### 2.7. Plasma protein binding models (PPBM) 1 and 2

Two plasma protein binding models (PPBM) were developed. PPBM 1 describes one ligand (either cortisol or cortisone) binding to two plasma proteins (CBG and albumin). PPBM 2 describes two ligands (both cortisol and cortisone) binding to both proteins. Thus, PPBM 2 can capture the competition between cortisol and cortisone for CBG and albumin binding, and the consequent effects on the unbound concentrations of both cortisol and cortisone. The one-ligand PPBM 1 treats cortisol and cortisone plasma protein binding as independent processes.

The full system of kinetic binding equations and ordinary differential equations for both PPBM 1 and 2 are provided in the Supplemental Materials (Section 1.3 and Supplemental Equations 1-19). Binding interactions are defined by association (k_on_) and dissociation (k_off_) rate constants, with the dissociation constant (K_d_) equal to k_off_ divided by k_on_. For modeling purposes, all k_on_ values were fixed at 100 μM⁻¹·s⁻¹, an upper bound based on a diffusion-limited on-rate (20). For each binding interaction, the corresponding k_off_ value is equal to the product of the k_on_ and the K_d_. Values of k_off_ were initially set to match literature K_d_ values (8,9,17), then varied to find the combination of parameters that best fit experimental observations. The ligand f_u_ was calculated as the ratio between the simulated unbound ligand concentration and the input total ligand concentration. Steady-state solutions of the governing differential equations (Supplemental Equations 1-8 for PPBM1, and Equations 9-19 for PPBM2) were found using the deterministic LSODA method implemented in COPASI (21) following a similar workflow as previously described (22). In this approach, model outputs are insensitive to the individual k_on_ and k_off_ values but sensitive to the K_d_ value. This was confirmed via sensitivity analysis varying k_on_ and k_off_ values but keeping the K_d_ constant.

### 2.8. COPASI implementation and parameter estimation

Parameter estimation was conducted using the Parameter Estimation task in COPASI. Model k_off_ values that minimized the objective function (the sum of squared residuals between simulated and observed unbound ligand concentrations) were estimated using the Levenberg–Marquardt nonlinear least-squares algorithm. PPBM 2 fits used a single objective function summing the residuals from both cortisol and cortisone measurements. The k_off_ values describing cortisol:albumin and cortisone:albumin binding were allowed to vary in the range of 5,000–100,000 s⁻¹ with initial values of 13,780 s⁻¹ and 44,000⁻¹ respectively (corresponding to reported K_d_ values of 138 µM (17) and 440 µM (9)). The k_off_ value for cortisol:CBG binding was allowed to vary between 0.01 and 20 s⁻¹ with an initial value of 3.3 s⁻¹ (corresponding to the literature K_d_ value of 0.033 μM (17)). The k_off_ value for cortisone:CBG binding was allowed to vary between 1 and 300 s⁻¹ with an initial value of 133 s⁻¹ (corresponding to the literature K_d_ value of 1.33 μM (8)). These parameter ranges were selected to encompass physiologically plausible binding affinities while allowing sufficient flexibility for optimization.

Following parameter estimation, observed and predicted unbound concentrations generated using COPASI were analyzed using MATLAB (MathWorks, Natick, MA). Prediction accuracy was evaluated using mean absolute percentage error (MAPE) and root mean squared error (RMSE). Agreement between predicted and observed concentrations was visualized using predicted-versus-observed plots including unity and two-fold error reference lines. In addition, the corrected Akaike Information Criterion (AIC) for model comparison was calculated according to equation 1 where n represents the number of observations (PPBM 1: n = 282; PPBM 2: n = 564), RSS represents log residual sum of squares, and k represents the number of estimated parameters (PPBM 1: k = 2; PPBM 2: k = 4).

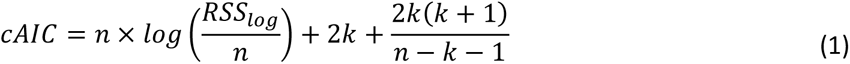

### 2.9. PPBM 2 bootstrap resampling

To evaluate the robustness and stability of the estimated k_off_ values in PPBM 2, a bootstrap resampling procedure was performed using the clinical study dataset. The original dataset consisted of 13 participants. For each bootstrap iteration, 10 participants were randomly selected without replacement while three participants were excluded (Supplemental Table 3). This 10x10 bootstrap design (10 resampled datasets each containing 10 participants) was implemented to assess the sensitivity of parameter estimates to participant level variability and to evaluate the consistency of model fitting across different resampled training sets. Bootstrap datasets were generated by resampling participants rather than individual observations to preserve the within participant structure of the data. For each selected participant, all observations from both study days and all time points were retained to maintain the correlation between repeated measurements and paired study conditions. For each bootstrap dataset, parameter estimation was repeated in COPASI using the same objective function and optimization settings described above. The resulting distribution of parameter estimates across bootstrap runs was used to quantify parameter uncertainty and model stability. Parameter estimates from the bootstrap runs were used to calculate 95% confidence intervals (CI) based on the mean and standard deviation of the estimates across the 10×10 bootstrap datasets, using a t-critical value of 2.262 (degrees of freedom = 9).

### 2.10. PPBM 2 sensitivity analysis

Sensitivity analysis of PPBM 2 was performed in MATLAB/Simulink to assess model performance and evaluate how physiological variability influences predicted unbound cortisol and cortisone concentrations. Baseline ligand concentrations were defined using the geometric mean values for albumin concentrations, and control day cortisol and cortisone C_max_ and CBG concentrations. The K_d_ values used were the final parameter estimates obtained from model fitting. In each simulation, these values were held constant unless explicitly varied as part of the sensitivity analysis.

Total cortisol and cortisone concentrations were varied from 0.02–2 μM. CBG concentrations were varied from 0.1–1 μM and albumin from 200–1,000 µM. The K_d_ values for cortisol:CBG and cortisone:CBG binding were varied from 0.001 to 0.3 μM and from 0.01 to 0.5 μM, respectively. The Kd values for cortisol:albumin and cortisone:albumin binding were varied from 50–1,000 μM and 100–2,000 μM, respectively. For each parameter set, the PPBM 2 was used to simulate unbound cortisol and cortisone concentrations.

### 2.11.

Cortisol and cortisone binding distributions in different physiological states

The final PPBM in MATLAB/Simulink was applied to simulate cortisol and cortisone binding across distinct physiological conditions in which CBG, albumin, cortisone, or cortisol concentrations are altered. These conditions included pregnancy, adrenal insufficiency, and liver disease and the PPBM 2 was implemented at steady state. Total cortisol, cortisone, CBG, and albumin concentrations were systematically varied across physiologically relevant ranges to predict unbound concentrations and f_u_ for both cortisol and cortisone in the different situations. The K_d_ values used were from the PPBM 2 fits, while ligand and binding protein concentrations were from reported literature values and are detailed in the Supplemental Materials Section 1.4. For healthy volunteers, model input concentrations were the average cortisol and cortisone Cmax values from the control day and literature reported values (23). Protein concentrations were from the control day of this study. The final total concentrations in the healthy population for cortisol, cortisone, CBG, and albumin were 0.310 µM, 0.056 µM, 0.431 µM, and 718 µM, respectively. To characterize ligand distribution under baseline conditions, total cortisol and cortisone concentrations were independently varied from 0–1 μM while holding all other inputs constant. Resulting concentrations and fractional distributions of unbound, CBG-bound, and albumin-bound cortisol and cortisone were evaluated at steady state to quantify changes in binding partitioning.

For pregnancy, values were derived from reported literature and represent trimester-dependent changes. Model inputs for total cortisol were 0.472, 0.712, 0.854, and 0.403 μM for first, second, and third trimester and postpartum, respectively; total cortisone inputs were 0.048, 0.072, 0.087, 0.041 μM; CBG concentrations were 0.700, 0.924, 1.140, and 0.574 μM; and albumin concentrations were 653, 573, 547, and 686 µM (Supplemental Table 4) (10,24–26).

For adrenal insufficiency, input values were determined from literature describing cortisol deficiency states (Supplemental Table 5) (27,28). Adrenal insufficiency simulations used cortisol 0.125 μM, cortisone 0.030 μM, CBG 0.548 μM, and albumin 541 µM.

For liver disease, input concentrations reflected literature reported values across increasing disease severity. In the reported studies, a reference population, and patients with liver disease classified as Child-Pugh’s class (CPC) A, B, and C, and cirrhosis were all included. The reference population cortisol, CBG, and albumin concentrations were 0.425 μM, 0.503 μM, and 618 μM, respectively. The reference population total cortisone concentrations were the same as the healthy population (0.056 μM). For CPC A cortisol, cortisone, CBG, and albumin concentrations were 0.587 μM, 0.050 μM, 0.574 μM, and 581 μM, respectively. For CPC B cortisol, cortisone, CBG, and albumin concentrations were 0.503 μM, 0.038 μM, 0.617 μM, and 510 μM, respectively. For CPC C, cortisol, cortisone, CBG, and albumin concentrations were 0.585 μM, 0.025 μM, 0.452 μM, and 451 μM, respectively. For cirrhosis, cortisol, cortisone, CBG, and albumin concentrations were 0.273 μM, 0.018 μM, 0.404 μM, 518 μM, respectively. All values and references are reported in Supplemental Table 6 (27,29–34).

### 2.12. Model and data availability

All model code, simulation workflows, and analysis scripts are publicly available in the following GitHub repository (https://github.com/Isoherranen-Lab/Cortisol-and-cortisone-plasma-protein-binding-model). The repository includes implementations of the plasma protein binding models (PPBM 1 and PPBM 2) in MATLAB/Simulink and COPASI, along with the corresponding model files. Code required to reproduce all figures presented in this study are also provided.

Additionally, the repository contains resources for model comparison, bootstrapping analyses, sensitivity analyses of PPBM 2, ligand distribution simulations, and simulations under varying physiological conditions. All results reported in this study can be reproduced using the provided code and model files.

## 3. Results

### 3.1. Cortisol and cortisone concentrations in healthy premenopausal women

The concentrations of cortisol and cortisone on the control study day and after a week of cortisol treatment are shown in Figure 1 and plots of individual participant data are shown in Supplemental Figures 1 and 2. The C_max_ values of cortisol and cortisone, along with average CBG and albumin concentrations, are summarized in Table 1. Both cortisol and cortisone concentrations increased significantly after seven days of treatment with hydrocortisone, synthetic cortisol (Table 1). The average f_u_ values for both cortisol and cortisone were significantly higher on the treatment day than on the control day (Table 1; Figure 2). The maximum change in cortisol f_u_ between control and treatment days was 1.9-fold, observed at the cortisol C_max_ after treatment. The maximum increase in cortisol f_u_ coincided with a 1.4-fold increase in cortisone f_u_. The increase in unbound peak concentrations (C_max,u_) was overall greater than the increase in total C_max_, 5.4- vs 2.5-fold for cortisol and 1.9- vs 1.3-fold for cortisone, respectively.

**Figure 1.**
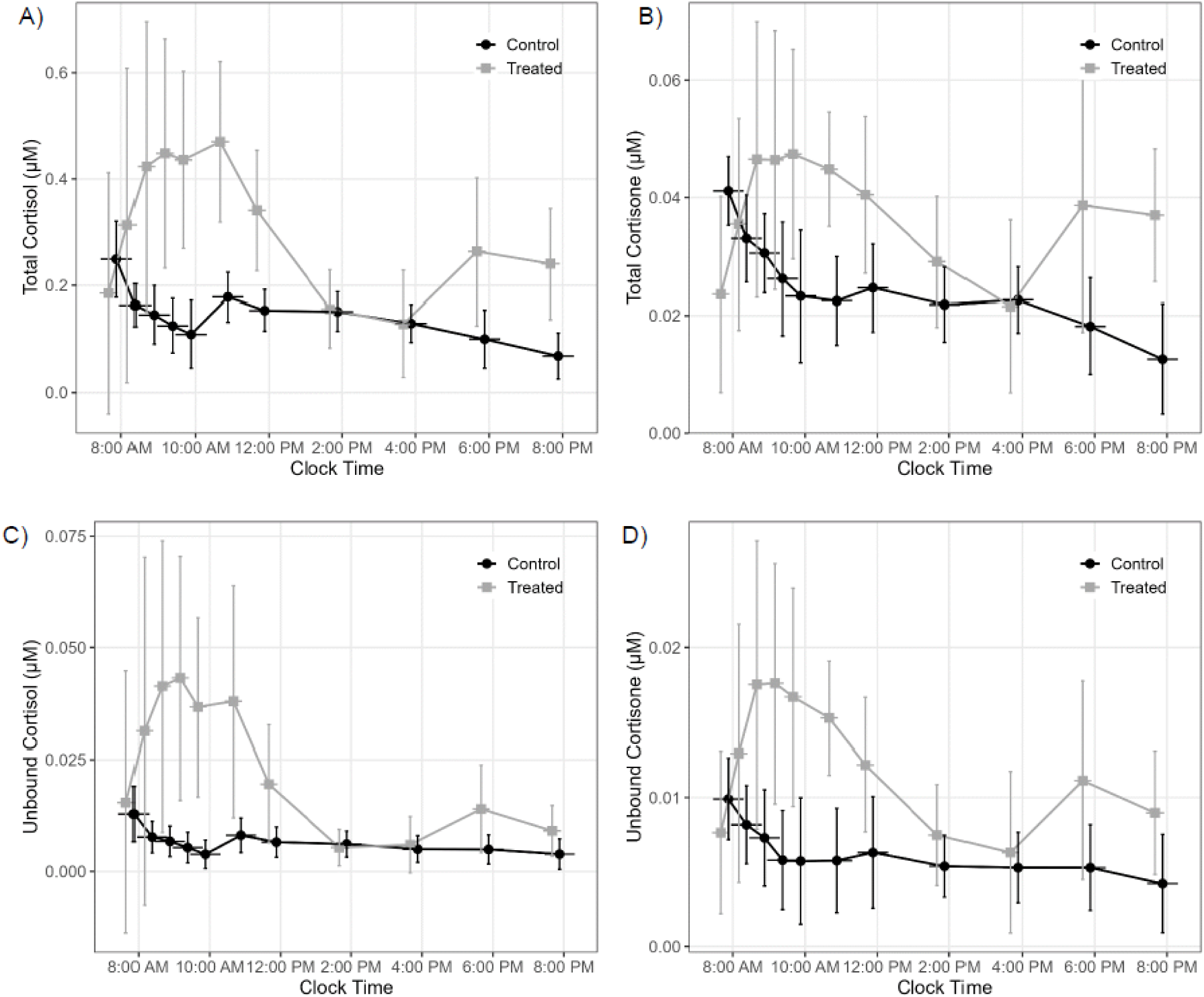
Cortisol and cortisone concentration-time curves on control and treated study days following 7 days of synthetic cortisol (hydrocortisone) administration. Total (A-B) and unbound (C-D) cortisol and cortisone are shown as a function of clock time. Data presented as average and standard deviation after excluding outliers. Outliers were identified using Grubbs’ test applied to the full dataset, as described in the statistical analysis section of the Materials and Methods.

**Figure 2.**
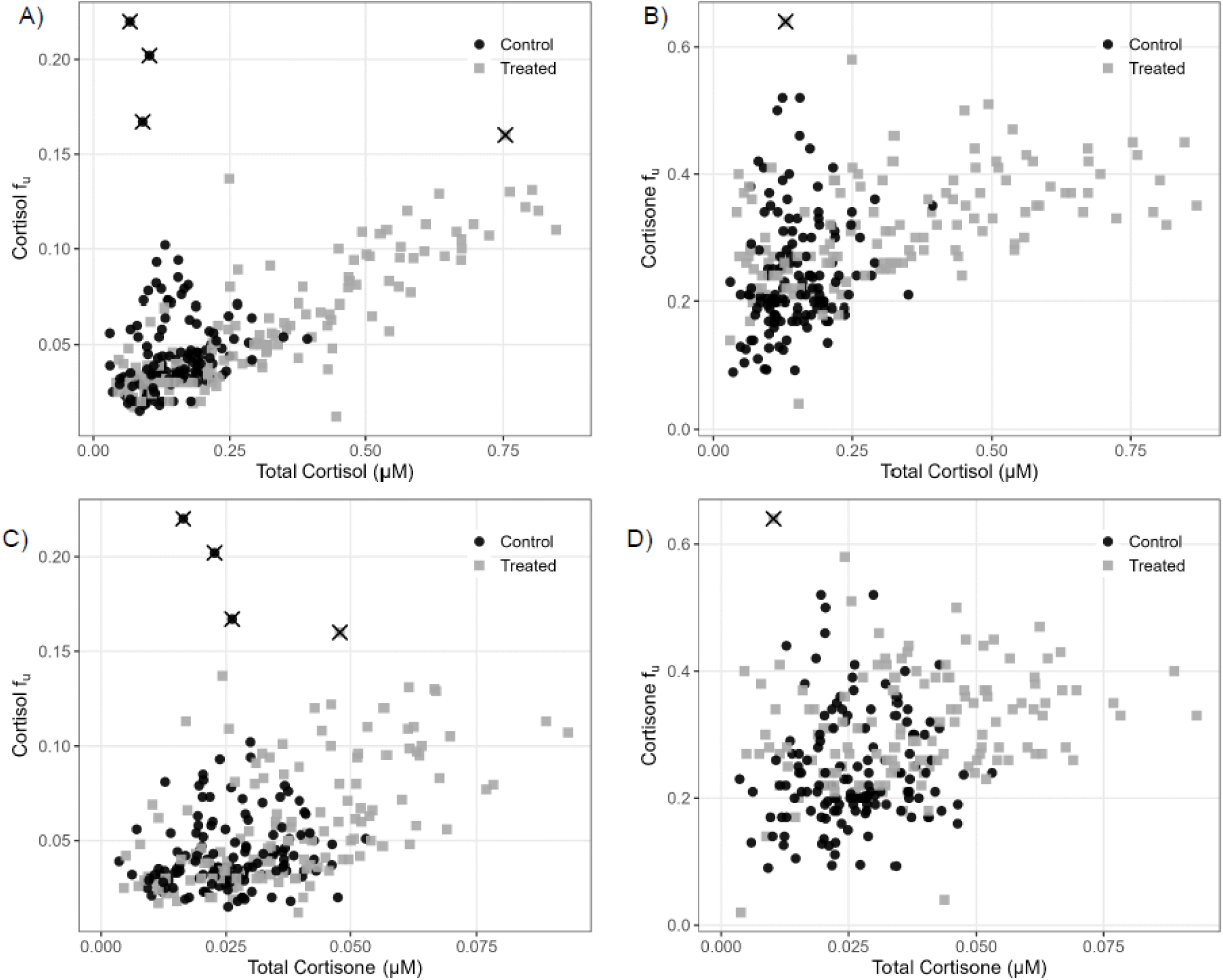
Relationship between total concentrations of cortisol (A-B) and cortisone (C-D) and fraction unbound (f_u_) of cortisol (A and C) and cortisone (B and D) on control (black circles) and treated (gray squares) study days. Outliers were identified using Grubbs’ test applied to the full dataset and presented as X symbols, as described in the statistical analysis section of the Materials and Methods.

**Table 1.** Cortisol, cortisone, corticosteroid-binding globulin (CBG), and albumin concentrations and binding in the clinical study on the control day and following seven days of treatment with synthetic, oral cortisol (hydrocortisone).

|  | Control | Treated | Treated/Control | p-value |
| --- | --- | --- | --- | --- |
| *Cortisol* |  |  |  |  |
| $C_{\max}$ ($\mu\text{M}$) | 0.257 (0.179-0.393) | 0.662 (0.471-0.869) | 2.5 (1.3-3.4) | 0.00000033 |
| $C_{\max,u}$ ($\mu\text{M}$) | 0.014 (0.0074-0.021) | 0.070 (0.039-0.121) | 5.4 (2.4-24) | 0.0000011 |
| Average $f_u$ | 0.040 (0.016-0.093) | 0.054 (0.38-0.064) | 1.4 (0.60-2.4) | 0.027 |
| Max $f_u$ | 0.062 (0.023-0.20) | 0.12 (0.083-0.16) | 1.9 (0.53-4.3) | 0.0038 |
| *Cortisone* |  |  |  |  |
| $C_{\max}$ ($\mu\text{M}$) | 0.042 (0.034-0.053) | 0.057 (0.035-0.093) | 1.3 (1.0-1.9) | 0.0011 |
| $C_{\max,u}$ ($\mu\text{M}$) | 0.011 (0.0083-0.018) | 0.021 (0.013-0.036) | 1.9 (1.1-4.0) | 0.00057 |
| Average $f_u$ | 0.23 (0.14-0.40) | 0.30 (0.23-0.39) | 1.3 (0.83-2.2) | 0.0045 |
| Max $f_u$ | 0.30 (0.22-0.52) | 0.43 (0.32-0.64) | 1.4 (0.85-2.0) | 0.0038 |
| *Plasma protein* |  |  |  |  |
| CBG ($\mu\text{M}$) | 0.431 (0.344-0.672) | 0.369 (0.292-0.505) | 0.82 (0.34-1.1) | 0.081 |
| Albumin ($\mu\text{M}$)<sup>‡</sup> | 718 (662-767) | | | |

Data presented as geometric mean and range; $C_{\max}$ : maximum concentration; $C_{\max,u}$ : unbound $C_{\max}$ ; Average $f_u$ : average fraction unbound in 12 h window; Max $f_u$ : maximum $f_u$ ; <sup>‡</sup>: measured on screening visit; statistical significance was defined as a paired t-test on log transformed data with FDR-adjusted p-values less than 0.05 and does not include outliers

Cortisol f_u_ was cortisol concentration-dependent with cortisol (Figure 2A), consistent with CBG saturation at higher cortisol concentrations. Cortisone f_u_ also increased with cortisol concentrations (Figure 2B). Cortisol f_u_ was not affected by increasing total cortisone concentrations (Figure 2C), whereas cortisone f_u_ showed a modest increase with higher cortisone concentrations (Figure 2D). This suggests that increased cortisol concentrations resulted in saturation of CBG binding and displacement of cortisone from CBG binding sites.

### 3.2. Comparison of PPBM 1 and 2

Fitting of the PPBMs produced consistent estimates of binding affinities across model structures (Table 2). Cortisol exhibited high affinity binding to CBG with estimated dissociation constants of 0.0141 μM in PPBM 1 and 0.0130 μM in PPBM 2. Cortisone binding to CBG was weaker, with estimated dissociation constants of 0.260 μM and 0.169 μM in PPBM 1 and 2, respectively. Binding to albumin was substantially lower affinity for both ligands, with estimated K_d_ values of approximately 173 or 172 µM for cortisol and 797 or 519 µM for cortisone in PPBM 1 and 2, respectively. The similarity of parameter estimates between PPBM 1 and PPBM 2 indicates stable identification of binding interactions despite differences in model structure.

**Table 2.**
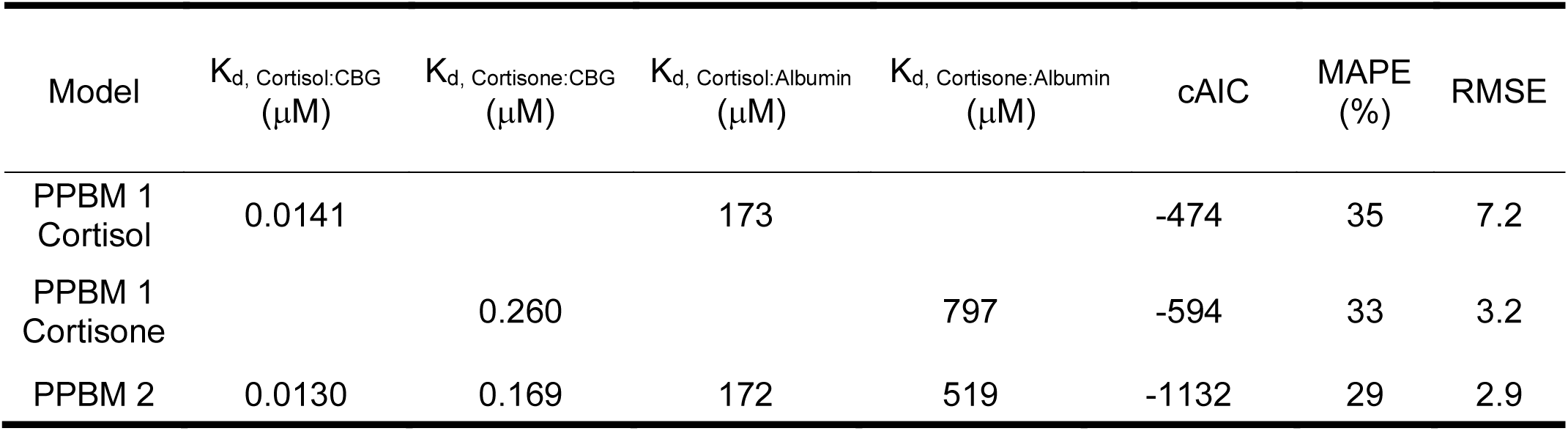
Estimated binding parameters for plasma protein binding models (PPBM 1 and PPBM 2), including dissociation constants (K_d_) for cortisol and cortisone interactions with corticosteroid-binding globulin (CBG) and albumin, along with corrected Akaike Information Criterion (cAIC), mean absolute percent error (MAPE), and root mean square error (RMSE).

Model predictive performance was evaluated using predicted-versus-observed unbound concentrations (Figure 3) and quantitative error metrics (Table 2). Across models, predictions demonstrated strong agreement with observed data (Figure 3). Model comparison using cAIC favored PPBM 2 (cAIC = −1132) over PPBM 1 for cortisol (−474) and cortisone (−594), indicating improved overall model support when both ligands were modeled simultaneously. Mean absolute percentage error was comparable between both PPBM 1 and PPBM 2 for both analytes (Table 2). Together, these results suggest that while PPBM 2 provides the preferred statistical description of the data, both models yield similar estimates of plasma protein binding affinities.

**Figure 3.**
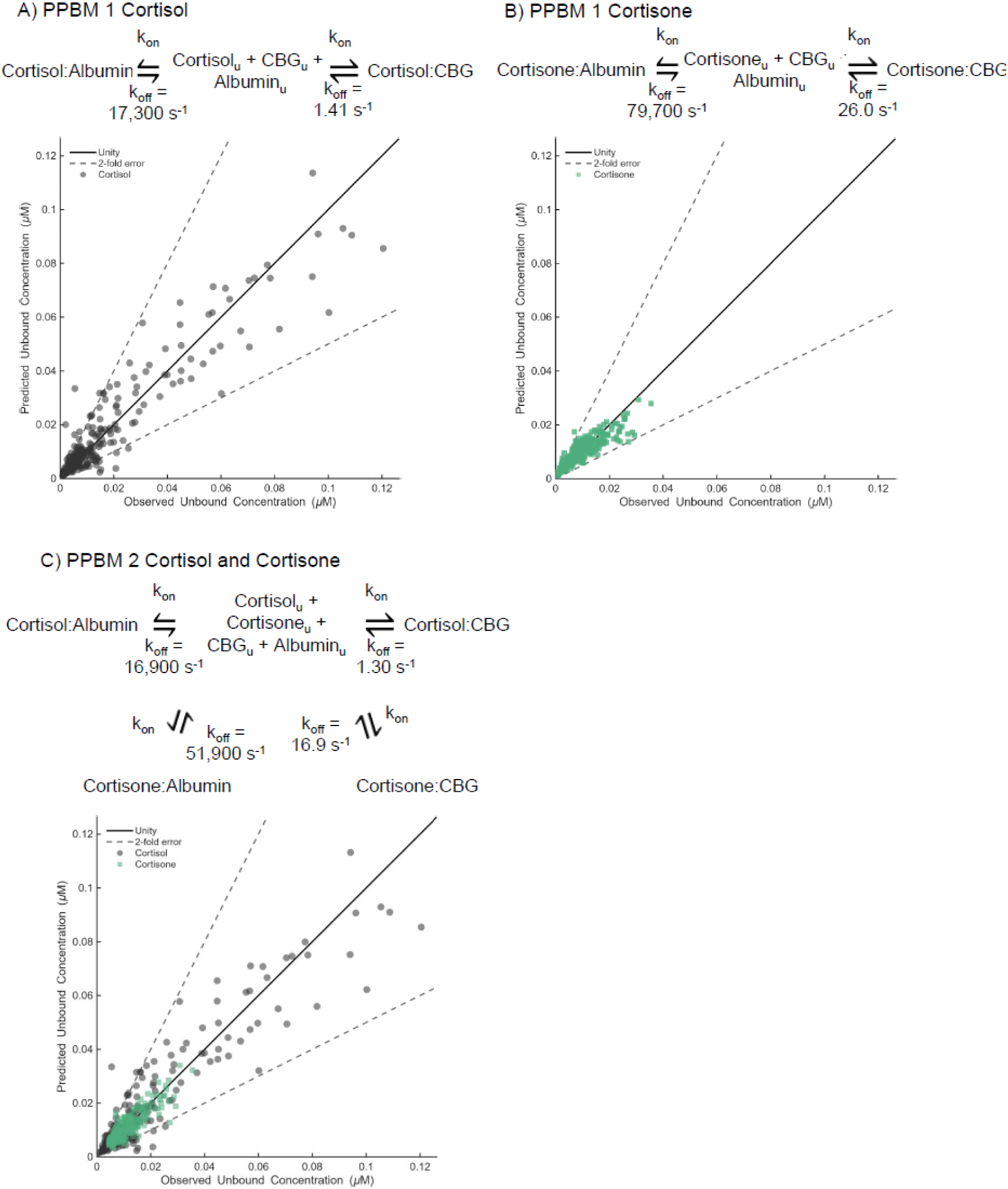
Unbound concentrations of cortisol and cortisone predicted using plasma protein binding models (PPBM) 1 and 2 versus observed concentrations. Panels A and B show results from PPBM 1, which incorporates protein binding by association (k_on_ = 100 µM⁻¹·s⁻¹) and dissociation (k_off_) rate constants. Panel A presents cortisol predictions, where 89% of simulated unbound fractions were within 2-fold of observed values. Panel B presents cortisone predictions with 95% of values within 2-fold error. Panel C shows results from PPBM 2, an integrated binding model describing both cortisol and cortisone simultaneously, yielding similarly strong agreement with observed data, with 89% of cortisol and 97% of cortisone predictions falling within 2-fold of observations.

### 3.3. Estimation of uncertainty in binding affinities

A bootstrap analysis was performed using repeated resampling to evaluate robustness of parameter estimation and predictive performance (Supplemental Table 7). Estimated K_d_ values remained stable across bootstrap replicates, indicating consistent identification of binding interactions. The resulting 95% confidence intervals (CI) of K_d_ values were 0.0077–0.0174 μM for cortisol:CBG binding and 0.133–0.178 μM for cortisone:CBG binding. Similarly, 95% CIs for albumin K_d_ values were 147–191 μM for cortisol:albumin and 95% CI of 397–647 μM for cortisone:albumin. The narrow parameter ranges relative to the magnitude of the parameters indicate stable estimation independent of participant resampling.

Predictive performance metrics were also consistent across bootstrap datasets, demonstrating that model accuracy was not sensitive to the specific composition of the cohort. Error metrics varied minimally across replicates, supporting reproducible prediction of unbound cortisol and cortisone concentrations (Supplemental Table 7). Together, these findings indicate that both parameter estimates and predictive capability of the PPBM are robust to variability within the clinical dataset.

### 3.4. PPBM 2 sensitivity analysis

PPBM 2 was implemented in MATLAB/Simulink for sensitivity analyses, exploring how predicted unbound cortisol and cortisone concentrations are influenced by variability in ligand and binding protein concentrations, and binding affinity parameters (Figure 4).

**Figure 4.**
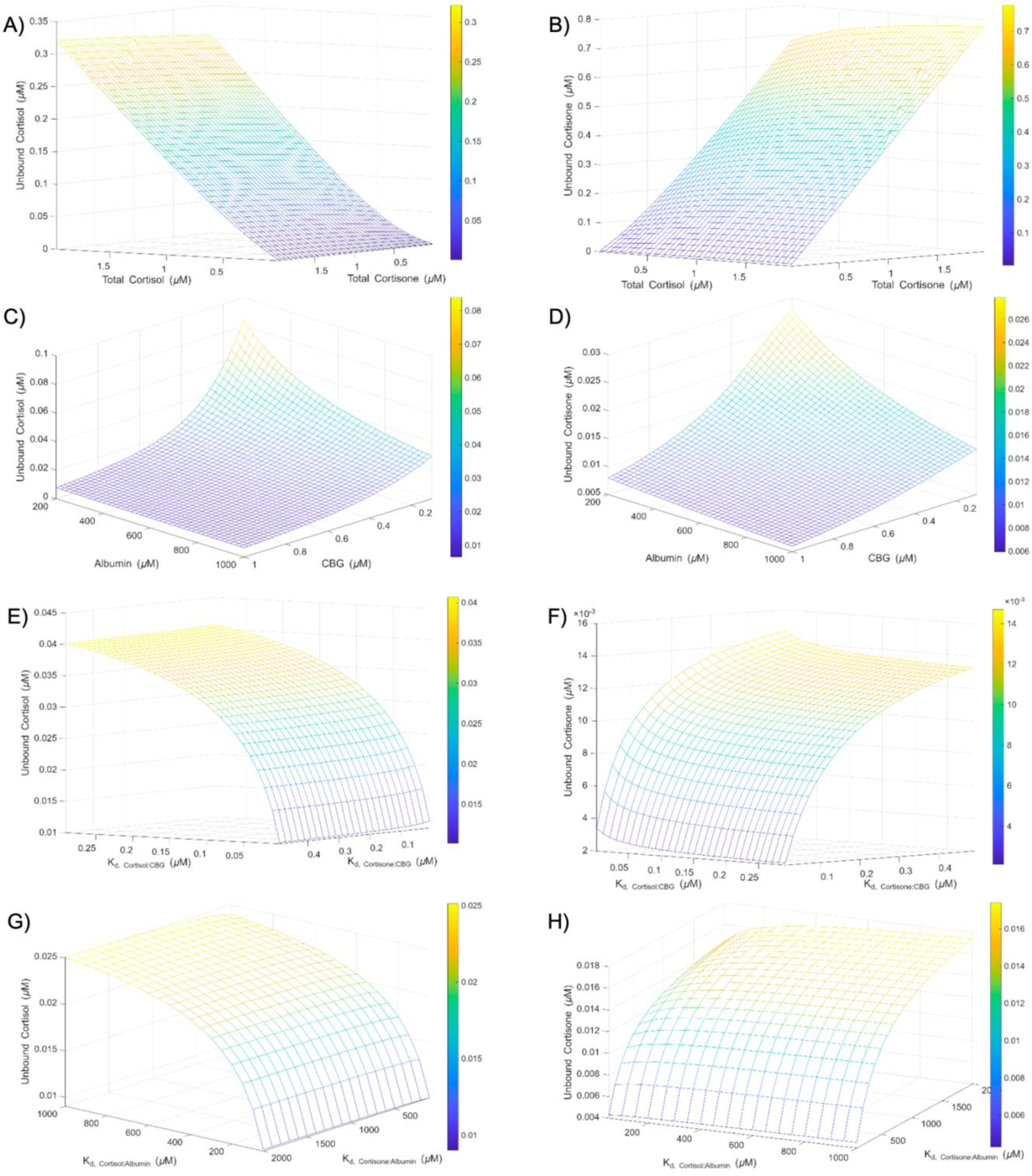
Sensitivity analysis of unbound cortisol (left column) and cortisone (right column) using PPBM 2. Sensitivity ranges were selected based on physiological concentrations, with fixed input values of total cortisol (0.310 μM), total cortisone (0.056 μM), total corticosteroid binding globulin (CBG; 0.431 μM), and total albumin (718 µM). The dissociation constants (K_d_) were from the optimized values. Panels A–B show the effect of varying total ligand concentrations, cortisol and cortisone. Panels C–D show the effect of varying protein concentrations, albumin and CBG. Panels E–F show the effect of varying K_d_ for cortisone:CBG and cortisol:CBG. Panels G-H show the effect of varying K_d_ for cortisone:albumin and cortisol:albumin.

The predicted unbound cortisol concentrations increased with increasing cortisol concentration but were minimally affected by increased total cortisone concentrations. Unbound cortisone concentrations did, however, increase with increasing total cortisone and cortisol concentrations, reflecting competitive displacement of cortisone from CBG binding sites by cortisol.

Predicted unbound concentrations of both cortisol and cortisone were sensitive to changes in CBG concentration. Decreasing CBG concentrations resulted in substantial increases in unbound cortisol and cortisone concentrations, highlighting the dominant role of CBG in cortisol and cortisone binding. Decreasing albumin concentrations resulted in increased unbound cortisol and cortisone concentrations with most pronounced effects observed when CBG concentrations were low. Under these conditions, binding of cortisol and cortisone to CBG is restricted and hence albumin binding and changes in albumin concentrations are more critical determinants of protein binding of cortisol and cortisone.

The unbound cortisol but not cortisone concentrations were sensitive to the cortisol:CBG K_d_. Conversely, unbound cortisone but not cortisol concentrations were sensitive to the cortisone:CBG K_d_. Similarly, unbound cortisol concentrations were sensitive to the cortisol: albumin K_d_ while unbound cortisone concentrations were sensitive to the cortisone:albumin K_d_. However, overall the unbound cortisol and cortisone concentrations were more sensitive to the K_d_ to CBG than to albumin binding.

### 3.5. Cortisol and cortisone binding distributions in different physiological states

The developed model was used to simulate the ligand distribution between unbound and protein bound states for cortisol and cortisone (Figure 5). At cortisol concentrations commonly observed in healthy adult population, the majority (>50%) of cortisol was predicted to be bound to CBG with 20-30% bound to albumin. With higher cortisol concentrations (> 0.9 μM) the fraction of cortisol bound to CBG decreased to ∼35% while the fraction bound to albumin was predicted to increase to ∼53%. Across the cortisol and cortisone concentrations considered, the unbound fraction of cortisol in plasma did not exceed 12% but increased by about 2-fold across the simulated cortisol concentration range (Figure 5). Overall increasing total cortisone or cortisol concentrations were predicted to decrease the fraction of cortisol and cortisone bound to CBG and increase the role of albumin in plasma protein binding of cortisol and cortisone. Cortisone had a higher unbound fraction (∼25%) when compared to cortisol (∼6%) across the concentration ranges and at typical cortisone concentrations a similar fraction of plasma cortisone was predicted to be bound to albumin and CBG.

**Figure 5.**
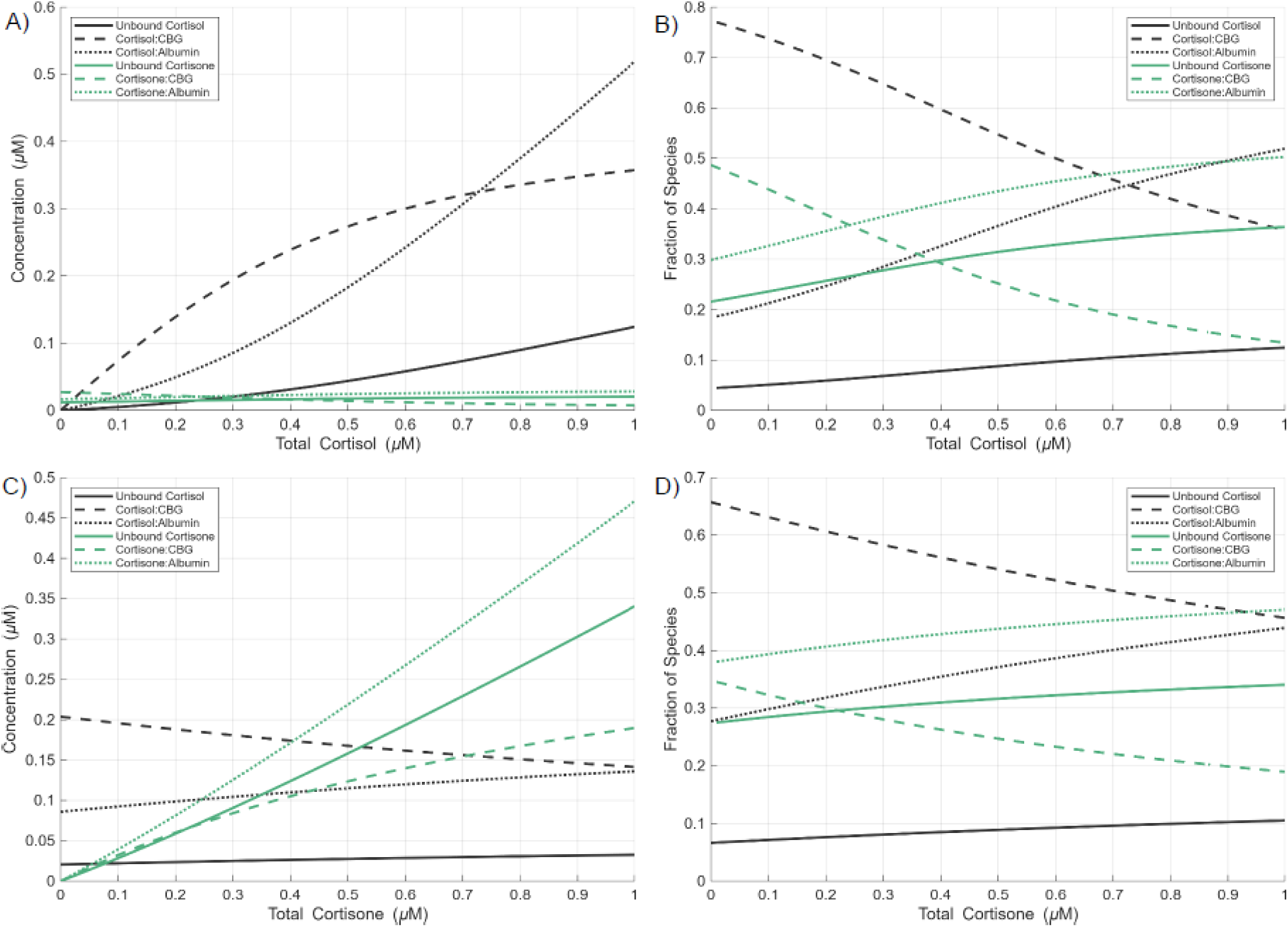
Simulated distribution of cortisol and cortisone species using the PPBM 2 across varying total cortisol (A,B) and cortisone (C,D) concentrations. Fixed input values corresponded to healthy reference population values for total cortisol (0.310 µM), total cortisone (0.056 µM), total corticosteroid-binding globulin (CBG; 0.431 µM), and total albumin (718 µM). The dissociation constants (K_d_) were set to the optimized values. Panels A–B show the effect of increasing total cortisol on (A) species concentrations and (B) fractional distribution. Panels C–D show the effect of increasing total cortisone on (C) species concentrations and (D) fractional distribution. Species include unbound (solid lines), CBG-bound (dashed lines), and albumin-bound (dotted-lines) cortisol (black) and cortisone (green).

The model application was expanded from healthy adults to predict the binding equilibria of cortisol and cortisone across multiple physiological states in which cortisol and cortisone concentrations are altered. These included pregnancy and postpartum, adrenal insufficiency, and liver disease (Figures 6-7). Model input protein and ligand concentrations were literature-reported physiological ranges (Supplemental Tables 4-6).

**Figure 6.**
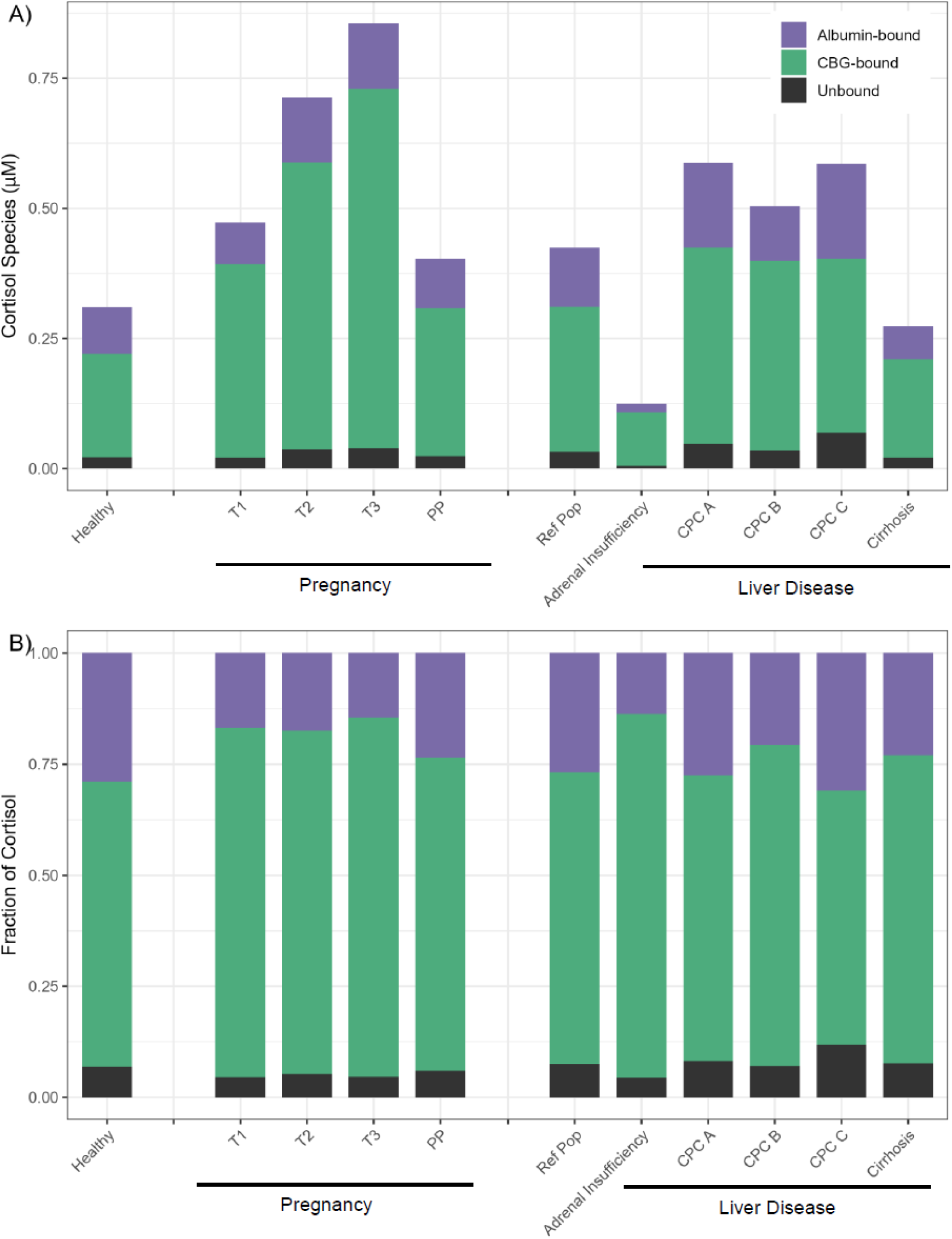
Cortisol concentrations (A) and fractions (B) bound to albumin, bound to corticosteroid-binding globulin (CBG), and unbound across different physiological states. States include a healthy reference population, pregnancy (trimester 1–3; T1–T3), postpartum (PP), adrenal insufficiency and liver disease (CPC: Child-Pugh’s Class). Simulations were generated using the PPBM 2 with final fitted K_d_ values. Ligand and plasma protein concentrations for each physiological state were assigned according to literature reported values specific to those conditions.

**Figure 7.**
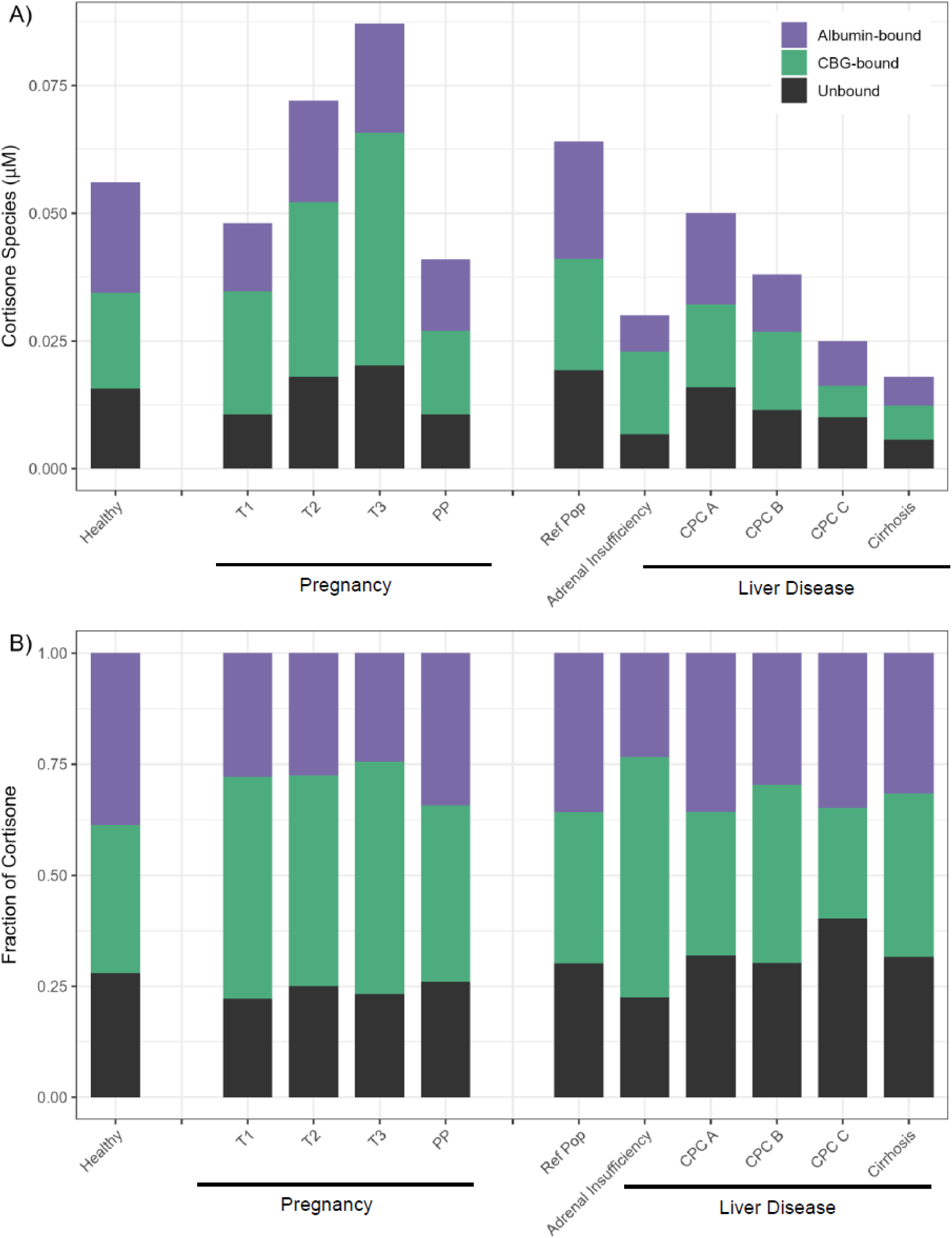
Cortisone concentrations (A) and fractions (B) bound to albumin, bound to corticosteroid-binding globulin (CBG), and unbound across different physiological states. States include a healthy reference population, pregnancy (trimester 1–3; T1–T3), postpartum (PP), adrenal insufficiency and liver disease (CPC: Child-Pugh’s Class). Simulations were generated using the PPBM 2 with final fitted K_d_ values. Ligand and plasma protein concentrations for each physiological state were assigned according to literature reported values specific to those conditions.

Across pregnancy, total cortisol, cortisone, and CBG concentrations increase and albumin concentrations decrease with gestational progression and return toward nonpregnant levels postpartum (Supplemental Table 4). Consequently, CBG binding is predicted to be dominant during pregnancy with cortisol and cortisone f_u_ predicted to decrease modestly in the third trimester. Due to the increased total cortisol and cortisone concentrations, unbound cortisol and cortisone concentrations were predicted to increase across pregnancy. Postpartum, binding patterns were similar to the healthy, nonpregnant reference population.

Adrenal insufficiency is characterized by reduced total cortisol and cortisone concentrations with minimal changes in binding protein levels relative to healthy individuals. Under these circumstances, binding distributions were predicted to be dominated by CBG binding. The unbound fractions of cortisol and cortisone were predicted to decrease considerably leading to greater decrease in unbound cortisol and cortisone concentrations than what would be expected from total concentrations alone (Figures 6-7).

With increasing liver disease severity, reported total cortisol and cortisone concentrations are greater in CPC A-C in comparison to cirrhosis (Supplemental Table 6). Both CBG and albumin concentrations decline relative to healthy individuals with more notable decrease in albumin. Model predictions, assuming unchanged cortisol synthesis, indicated a progressive increase in unbound cortisol and cortisone concentrations and f_u_ from CPC A to CPC C (Figures 6–7). CBG-bound fractions of cortisol and cortisone decreased from CPC B to C, with a corresponding increase in albumin binding, reflecting a shift toward lower-affinity binding. In cirrhosis, the significant decrease in cortisol concentrations together with marked reduction in albumin binding capacity were predicted to result in considerably reduced free cortisol and cortisone.

## 4. Discussion

Accurate interpretation of plasma cortisol concentrations is limited by nonlinear plasma protein binding and physiological variability in both ligand and binding protein concentrations. Although total cortisol concentrations are routinely measured, only unbound cortisol is available for receptor activation, metabolism, and elimination (2,22,23). This distinction becomes particularly important across physiological and disease states in which CBG, albumin, cortisol, and cortisone concentrations are substantially altered resulting in altered unbound fraction of cortisol. Presence of cortisone further complicates interpretation of cortisol concentrations as cortisone also binds to CBG and is reversibly converted to cortisol by 11β-hydroxysteroid dehydrogenases (24). Consequently, mechanistic interpretation of cortisol disposition requires quantitative description of how cortisol and cortisone partition among unbound, CBG-bound, and albumin-bound species under varying physiological conditions.

The present study developed a plasma protein binding model (PPBM) that quantitatively describes binding of cortisol and cortisone to CBG and albumin under physiological conditions. By integrating both ligands and both binding proteins into a single model, the PPBM enables simultaneous prediction of unbound cortisol and cortisone concentrations and the binding states of both ligands across a wide range of physiological states. Model parameters were estimated using *in vivo* data from healthy volunteers, with wide range of cortisol and cortisone concentrations. The cortisol:CBG and cortisol:albumin K_d_ values (0.0130 μM and 172 μM, respectively) determined here were similar to the previously published K_d_ values estimated from *in vivo* plasma protein binding (0.01 μM (35) and 138 μM (17), respectively). In contrast, the K_d_ values for CBG binding measured *in vitro* using recombinant glycosylated human CBG and surface plasmon resonance (SPR) for cortisol (0.033-0.21 μM) and for cortisone (1.34 μM) (8) were 3-10 fold higher than the binding affinities estimated here. For cortisone albumin binding, our fitted value (519 μM) was similar to the previously estimated K_d_ measured by equilibrium dialysis in buffer with human serum albumin (440 μM (9)). These comparisons suggest that the protein environment in plasma and the physiological context may affect cortisol and cortisone binding with CBG but not albumin. This further indicates that *in vitro* assays do not adequately capture the binding kinetics observed in plasma. As such, the modeling approach adopted here to explore CBG binding affinities using unbound fractions determined in plasma samples is particularly valuable and may be applied to other physiological conditions to explore potential changes in CBG binding behavior.

Differences between *in vitro* and *in vivo* K_d_ estimates for CBG binding likely reflect both assay specific limitations and underlying biological complexity. *In vitro* measurements are highly dependent on experimental conditions, including protein and ligand concentration ranges, buffer composition, and ionic environment, all of which can influence protein conformation and electrostatic interactions (36–38). In particular, buffer components may interact with protein surfaces or alter hydration states, thereby affecting binding behavior (37,38). Additionally, physicochemical factors such as pH and temperature can influence protein structure and binding kinetics, and may not be consistently replicated across *in vitro* studies (8). This is particularly relevant in the case of CBG, whose affinity for steroid ligands is highly temperature-sensitive (8,39). In contrast, plasma samples represent a multi-protein system in which ligands distribute across binding partners with differing affinities and capacities and in which protein-protein interactions and an *in vivo*-like crowded protein environment can be replicated. While albumin binding did not appear sensitive to such potential factors, our data suggest that for CBG binding kinetics, significant discrepancies between *in vitro* and *in vivo* binding estimates exist and protein environment has a significant impact on ligand binding kinetics. As such, the use of data integrated approaches to capture physiologically relevant binding behavior is recommended.

Model performance suggests that the observed differences between *in vitro* and *in vivo* K_d_ values are unlikely to arise from instability or overfitting of the PPBM. Strong agreement between predicted and observed unbound cortisol and cortisone concentrations (Figure 3) indicates that the model accurately captures binding behavior across the studied concentration ranges and between individual study participants. In addition, comparable binding affinities obtained from structurally distinct PPBMs suggest that parameter estimates are not dependent on a specific model structure, supporting identifiability of the underlying binding interactions. Bootstrap analyses further demonstrated that parameter estimation and predictive performance were not sensitive to the composition of the clinical dataset, indicating robustness to data resampling. This provides confidence that the fitted K_d_ values are consistent and not model-specific artifacts and reinforces that discrepancies with *in vitro* estimates are more likely attributable to physiological complexity and experimental differences than to limitations in model performance.

Sensitivity analyses provided insight into specific impact of both ligand and protein interactions on unbound cortisol and cortisone concentrations. Predicted unbound cortisol concentrations were most strongly influenced by total cortisol and CBG concentrations and cortisol:CBG binding affinity, highlighting the dominant regulatory role of high-affinity CBG interactions. Albumin exerted a secondary buffering effect that became more apparent when CBG concentrations were decreased, consistent with the lower-affinity, higher-capacity binding characteristics of albumin (6,29). The sensitivity analyses (Figure 4) and ligand species distribution (Figure 5) demonstrate that cortisol is predominantly bound to CBG with a smaller fraction bound to albumin, consistent with previous reports of ∼5% unbound, ∼80% CBG-bound, and ∼15% albumin-bound *in vivo* (5,6,8).

To our knowledge, this is the first study to characterize cortisone distribution across plasma protein binding partners. Our analysis demonstrates that cortisone undergoes similar redistribution as cortisol driven by cortisol-mediated displacement of cortisone from CBG. Competitive interactions between cortisol and cortisone were reflected in reciprocal effects on unbound concentrations, with increases in cortisol concentrations resulting in displacement of cortisone from CBG (Figures 4-5). These findings confirm that cortisone binding behavior is governed primarily by CBG-mediated saturation dynamics rather than by albumin availability within the tested physiological concentration ranges.

The PPBM was successfully extrapolated to simulate cortisol and cortisone binding states across physiological conditions. During pregnancy, progressive increases in total cortisol and CBG concentrations were predicted to produce limited effects in unbound fraction but substantial increases in unbound cortisol concentrations across trimesters when compared to postpartum. These predictions align with reported pregnancy-associated changes in cortisol plasma concentrations and illustrate how coordinated changes in ligand and binding protein concentrations shift binding equilibria (10). In adrenal insufficiency, where binding protein levels are preserved, reduced ligand concentrations led to lower predicted unbound fractions of cortisol. In liver disease, decreased CBG and albumin concentrations were predicted to increase unbound fractions of cortisol and cortisone as disease severity progresses from CPC A to C. However, in cirrhosis, unbound fractions were predicted to decrease compared to earlier liver disease stages. Overall, these simulations assume that binding affinities remain constant across physiological conditions, an assumption that should be evaluated in future studies. Apparent dissociation constants may vary due to changes in CBG glycosylation, protein heterogeneity, or ligand competition (6,8,14). The modeling methodology presented here can be applied in the future to estimate how ligand binding affinity to CBG may be altered in specific populations and physiological states.

In conclusion, the PPBM provides a quantitative framework for describing nonlinear and competitive binding of cortisol and cortisone *in vivo* and for predicting unbound plasma concentrations across physiological states. By enabling kinetic interpretation of how binding protein dynamics regulate unbound cortisol, this work advances understanding of endocrine regulation and underscores the limitations of total hormone measurements. The framework supports improved interpretation of endocrine biomarkers, can be incorporated into pharmacokinetic and physiologically based pharmacokinetic models, and may be applied to evaluate other endogenous or xenobiotic disposition under changing physiological or pathological states.

## Supporting information

Biorender license

## Abbreviations

11β-HSD: 11β-hydroxysteroid dehydrogenase
AIC: Akaike Information Criterion
CAD: collision activated dissociation
CBG: corticosteroid binding globulin
CE: collision energy
CXP: collision cell exit potential
DP: declustering potential
ELISA: enzyme linked immunosorbent assay
EP: entrance potential
FDR: false discovery rate
f_u_: fraction unbound
GS1/2: source gas 1/2
HQC: high quality control
IRB: institutional review board
KPi: potassium phosphate buffer
K_d_: dissociation constant
k_off_: off rate
k_on_: on rate
LC-MS/MS: liquid chromatography-tandem mass spectrometry
LQC: low quality control
MAPE: mean absolute percent error
MQC: middle quality control
MRM: multiple reaction monitoring
MPE: mean percent error
PPBM: plasma protein binding model
QC: quality control
RED: rapid equilibrium dialysis

## Data availability

The authors declare that all of the data supporting the finding of this study are available within the paper, supplemental materials, and at the following GitHub repository (https://github.com/Isoherranen-Lab/Cortisol-and-cortisone-plasma-protein-binding-model).

## CRediT authorship contribution statement

Aurora Authement: Writing — original draft, Visualization, Methodology, Investigation, Formal analysis, Funding acquisition, Conceptualization.

Abhinav Nath: Writing — review and editing, Formal analysis

Katya B Rubinow: Writing — review and editing, Investigation, Supervision, Project administration

John K Amory: Writing — review and editing, Investigation, Supervision, Project administration

Nina Isoherranen: Writing — review and editing, Conceptualization, Supervision, Project administration, Methodology, Investigation, Funding acquisition.

