## Supplemental materials for "Kinetics of cortisol and cortisone binding to corticosteroid binding globulin and albumin *in vivo*"

#### 1. Supplemental Materials and Methods

##### *1.1 Healthy volunteer clinical study sampling time*

Supplemental Table 1. The clock time of dronabinol dosing on each clinical study day (control and cortisol treated). The blood draw times are counted from this clock time being time 0.

| Participant ID | Control Day | Treated Day |
|---|---|---|
| 1 | 7:45 AM | 7:21 AM |
| 2 | 8:40 AM | 7:30 AM |
| 5 | 7:50 AM | 7:50 AM |
| 6 | 7:55 AM | 7:55 AM |
| 8 | 8:20 AM | 8:00 AM |
| 10 | 7:40 AM | 7:40 AM |
| 12 | 7:30 AM | 7:45 AM |
| 13 | 8:00 AM | 7:25 AM |
| 15 | 8:35 AM | 7:40 AM |
| 17 | 7:35 AM | 8:05 AM |
| 18 | 7:50 AM | 7:35 AM |
| 19 | 7:15 AM | 7:20 AM |
| 21 | 7:30 AM | 7:35 AM |
| Geometric Mean | 7:52 AM | 7:40 AM |
| Minimum | 7:15 AM | 7:20 AM |
| Maximum | 8:40 AM | 8:05 AM |

*1.2. Quantification of cortisol and cortisone in plasma via LC-MS/MS*

Supplemental Table 2. Method validation data for plasma analysis. Accuracy and precision for cortisol and cortisone at nominal concentrations in charcoal stripped human serum.

| | LLQC | | LQC | | MQC1 | | MQC2 | | HQC | |
|---|---|---|---|---|---|---|---|---|---|---|
| | %Error | %CV | %Error | %CV | %Error | %CV | %Error | %CV | %Error | %CV |
| *Cortisol* | 25 nM | | 100 nM | | 350 nM | | 500 nM | | 750 nM | |
| Inter-injection | -1.5 | 2.8 | -1.1 | 3.4 | -0.53 | 3.8 | -4.2 | 3.1 | 0.67 | 1.8 |
| Intraday | 11 | 3.9 | 4.8 | 9.2 | 0.31 | 7.4 | 8.5 | 4.4 | -1.1 | 4.6 |
| Interday | -1.2 | 7.5 | -2.9 | 4.1 | 1.6 | 1.4 | 13 | 2.7 | -1.4 | 2.3 |
| 48 h Stability | -5.6 | 6.0 | -3.4 | 5.4 | 1.3 | 3.0 | 11 | 1.8 | -4.9 | 5.2 |
| *Cortisone* | 15 nM | | 25 nM | | 30 nM | | 40 nM | | 60 nM | |
| Inter-injection | -1.8 | 1.4 | 3.6 | 2.9 | 4.1 | 2.4 | -0.89 | 1.8 | 1.7 | 2.1 |
| Intraday | 9.2 | 5.8 | 1.6 | 10 | -7.2 | 4.7 | -8.7 | 6.5 | 4.7 | 2.9 |
| Interday | 5.8 | 9.1 | -1.2 | 8.1 | -3.1 | 5.9 | -2.2 | 5.8 | -1.1 | 6.5 |
| 48 h Stability | 6.7 | 8.8 | 4.6 | 7.8 | -7.2 | 4.7 | -3.8 | 5.6 | -4.7 | 7.5 |

%Error: (reference-measured)/measured; %CV: coefficient of variation; LLQC: lower limit of quantification QC; LQC: low QC; MQC1 or 2: middle QC 1 or 2; HQC: high QC; NA: not applicable

*1.3. Plasma protein binding models (PPBM) ordinary differential equations*

PPBM 1 is governed by the following system of mass balance and differential equations, where *CS* represents either cortisol or cortisone:

$$[CS]_t = [CS]_u + [CS{:}\, CBG] + [CS{:}\, Albumin] \quad (1)$$

$$[CBG]_t = [CBG]_u + [CS{:}\, CBG] \quad (2)$$

$$[Albumin]_t = [Albumin]_u + [CS{:}\, Albumin] \quad (3)$$

$$\frac{d[CS{:}CBG]}{dt} = k_{on,CS:CBG}[CS]_u[CBG]_u - k_{off,CS:CBG}[CS{:}\, CBG] \quad (4)$$

$$\frac{d[CS{:}Albumin]}{dt} = k_{on,CS:Albumin}[CS]_u[Albumin]_u - k_{off,CS:Albumin}[CS{:}\, Albumin] \quad (5)$$

$$\frac{d[CS]_u}{dt} = k_{off,CS:CBG}[CS{:}\, CBG] - k_{on,CS:CBG}[CS]_u[CBG]_u + k_{off,CS:Albumin}[CS{:}\, Albumin] - k_{on,CS:Albumin}[CS]_u[Albumin]_u \quad (6)$$

$$\frac{d[CBG]_u}{dt} = k_{off,CS:CBG}[CS{:}\, CBG] - k_{on,CS:CBG}[CS]_u[CBG]_u \quad (7)$$

$$\frac{d[Albumin]_u}{dt} = k_{off,CS:Albumin}[CS{:}\, Albumin] - k_{on,CS:Albumin}[CS]_u[Albumin]_u \quad (8)$$

CS represents either cortisol or cortisone and will be referred to as ligand. Here, $[CS]_u$ represents unbound ligand, $[CBG]_u$ is unbound CBG, $[Albumin]_u$ is unbound albumin. The [CS:CBG] is ligand bound to CBG concentration, and [CS:Albumin] is ligand bound to albumin concentration. The binding kinetics are described in detail below.

PPBM 2 is governed by the following system of equations:

$$[Cortisol]_t = [Cortisol]_u + [Cortisol\!:CBG] + [Cortisol\!:Albumin] \quad (9)$$

$$[Cortisone]_t = [Cortisone]_u + [Cortisone\!:CBG] + [Cortisone\!:Albumin] \quad (10)$$

$$[CBG]_t = [CBG]_u + [Cortisol\!:CBG] + [Cortisone\!:CBG] \quad (11)$$

$$[Albumin]_t = [Albumin]_u + [Cortisol\!:Albumin] + [Cortisone\!:Albumin] \quad (12)$$

$$\frac{d[Cortisol\!:CBG]}{dt} = k_{on,Cortisol:CBG}[Cortisol]_u[CBG]_u - k_{off,Cortisol:CBG}[Cortisol\!:CBG] \quad (13)$$

$$\frac{d[Cortisol\!:Albumin]}{dt} = k_{on,Cortisol:Albumin}[Cortisol]_u[Albumin]_u - k_{off,Cortisol:Albumin}[Cortisol\!:Albumin] \quad (14)$$

$$\frac{d[Cortisone\!:CBG]}{dt} = k_{on,Cortisone:CBG} \times [Cortisone]_u[CBG]_u - k_{off,Cortisone:CBG}[Cortisone\!:CBG] \quad (15)$$

$$\frac{d[Cortisol]_u}{dt} = k_{off,Cortisol:CBG}[Cortisol\!:CBG] - k_{on,Cortisol:CBG}[Cortisol]_u[CBG]_u + k_{off,Cortisol:Albumin}[Cortisol\!:Albumin] - k_{on,Cortisol:Albumin}[Cortisol]_u[Albumin]_u \quad (16)$$

$$\frac{d[Cortisone]_u}{dt} = k_{off,Cortisone:CBG}[Cortisone\!:CBG] - k_{on,Cortisone:CBG}[Cortisone]_u[CBG]_u + k_{off,Cortisone:Albumin}[Cortisone\!:Albumin] - k_{on,Cortisone:Albumin}[Cortisone]_u[Albumin]_u \quad (17)$$

$$\frac{d[CBG]_u}{dt} = k_{off,Cortisol:CBG}[Cortisol\!:CBG] - k_{on,Cortisol:CBG}[Cortisol]_u[CBG]_u + k_{off,Cortisone:CBG}[Cortisone\!:CBG] - k_{on,Cortisone:CBG}[Cortisone]_u[CBG]_u \quad (18)$$

$$\frac{d[Albumin]_u}{dt} = k_{off,Cortisol:Albumin}[Cortisol: Albumin] - k_{on,Cortisol:Albumin}[Cortisol]_u[Albumin]_u + k_{off,Cortisone:Albumin}[Cortisone: Albumin] - k_{on,Cortisone:Albumin}[Cortisone]_u[Albumin]_u \quad (19)$$

Here, $[Cortisol]_u$ is unbound cortisol concentration, [Cortisol:CBG] is cortisol bound to CBG concentration, and [Cortisol:Albumin] is cortisol bound to albumin concentration. The $[Cortisone]_u$ is unbound cortisone concentration, and [Cortisone:CBG] is cortisone bound to CBG concentration. $[CBG]_t$ is the total CBG concentration, $[CBG]_u$ is unbound CBG concentration, and [Cortisol:CBG] and [Cortisone:CBG] represent CBG bound to cortisol and cortisone concentrations. $[Albumin]_t$ is the total albumin concentration, $[Albumin]_u$ is unbound albumin concentration, and [Cortisol:Albumin] is cortisol bound to albumin concentration.

Binding interactions are defined by association ($k_{on}$) and dissociation ($k_{off}$) rate constants, with the dissociation constant ($K_d$) equal to $k_{off}$ divided by $k_{on}$. For modeling purposes, all $k_{on}$ values were fixed at 100 $\mu M^{-1} \cdot s^{-1}$, an upper bound based on the diffusion-limited on-rate (1). For each binding interaction, the corresponding $k_{off}$ value is equal to the product of the $k_{on}$ and the $K_d$. Values of $k_{off}$ were initially set to match literature $K_d$ values (2–4), then varied to find the combination of parameters that best fit experimental observations.

Supplemental Table 3. Participant identification (SID) assignment to training and test datasets, with 10 participants in each training set.

| Split | Train SID | Test SID |
|---|---|---|
| 1 | 1,6,7,11,3,12,2,13,5,4 | 8,10,9 |
| 2 | 5,10,8,11,2,4,3,9,13,1 | 6,7,12 |
| 3 | 8,11,3,1,6,13,4,9,10,2 | 7,5,12 |
| 4 | 5,4,2,12,3,1,11,7,9,6 | 13,8,10 |
| 5 | 2,12,9,13,11,1,3,10,5,8 | 4,7,6 |
| 6 | 1,9,5,4,10,12,2,8,13,7 | 6,11,3 |
| 7 | 1,8,13,2,3,12,4,6,11,7 | 9,10,5 |
| 8 | 8,4,5,2,6,13,1,9,11,10 | 12,7,3 |
| 9 | 3,5,2,8,10,6,13,9,4,11 | 12,7,1 |
| 10 | 13,12,9,1,8,10,6,11,3,5 | 7,4,2 |

*1.4. Literature references for plasma protein binding model parameter inputs for different physiological conditions – pregnancy, adrenal insufficiency, and liver disease*

Supplemental Table 4. Reported values for cortisol, cortisone, corticosteroid binding globulin (CBG), and albumin concentrations during first, second, and third trimesters (T1-T3) and postpartum (PP). Only literature values that included postpartum comparison were included in the analysis.

| | T1 | T2 | T3 | PP | Reference |
|---|---|---|---|---|---|
| ***Cortisol (μM)*** | | | | | |
| | 0.577 | 0.878 | 1.043 | 0.486 | (5) |
| | 0.386 | 0.790 | 0.792 | 0.296 | (6) |
| | | | 0.910 | 0.441 | (7) |
| | | 0.520 | 0.708 | 0.389 | (8) |
| GM | 0.472 | 0.712 | 0.854 | 0.403 | |
| ***Cortisone (μM)*** | | | | | |
| | | | 0.116 | 0.025 | (7) |
| Calculated | 0.048 | 0.072 | 0.087 | 0.41 | |
| **CBG *(μM)*** | | | | | |
| | 0.700 | 0.920 | 1.107 | 0.657 | (5) |
| | | 0.929 | 1.174 | 0.502 | (8) |
| GM | 0.700 | 0.924 | 1.140 | 0.574 | |
| ***Albumin (μM)*** | | | | | |
| | 653 | 573 | 547 | 686 | (9) |

GM: geometric mean; Calculated: calculated ratios of cortisone/cortisol from T3 and PP and multiplied by the geometric mean of cortisol

One study that included cortisol and CBG concentrations was excluded due to CBG concentrations being ~2-fold greater than the geometric mean of the two included studies (10).

Supplemental Table 5. Literature reported values for cortisol, cortisone, corticosteroid binding globulin (CBG), and albumin concentrations during adrenal insufficiency.

| | Adrenal Insufficiency | Reference |
|---|---|---|
| ***Cortisol (μM)*** | | |
| | 0.061 | (11) |
| | 0.188 | (11) |
| | 0.110 | (12) |
| GM | 0.125 | |
| ***Cortisone (μM)*** | | |
| | 0.010-0.050 | (12) |
| Midpoint | 0.030 | |
| ***CBG (μM)*** | | |
| | 0.538 | (11) |
| | 0.558 | (11) |
| | 0.450 | (12) |
| GM | 0.548 | |
| ***Albumin (μM)*** | | |
| | 556 | (11) |
| | 526 | (11) |
| | 630 | (12) |
| GM | 541 | |

GM: geometric mean

Supplemental Table 6. Literature reported values for cortisol, cortisone, corticosteroid binding globulin (CBG), and albumin concentrations during liver disease and cirrhosis.

| Reference | Ref Pop | CPC A | CPC B | CPC C | Cirrhosis | Reference |
|---|---|---|---|---|---|---|
| ***Cortisol (μM)*** | | | | | | |
| | | 0.800 | 0.690 | 0.585 | | (13) |
| | 0.328 | 0.490 | | | | (14) |
| | 0.736 | 0.723 | | | | (11) |
| | 0.317 | | | | 0.273 | (15) |
| | | 0.420 | 0.367 | | | (16) |
| GM | 0.425 | 0.587 | 0.503 | 0.585 | 0.273 | |
| ***Cortisone (μM)*** | | | | | | |
| | | 0.020-0.080 | 0.015-0.060 | 0.010-0.040 | 0.005-0.030 | (17) |
| Midpoint | | 0.050 | 0.0375 | 0.025 | 0.0175 | |
| ***CBG (μM)*** | | | | | | |
| | 0.4552 | 0.474 | | | | (14) |
| | 0.519 | 0.538 | | | | (11) |
| | 0.538 | | | | 0.404 | (15) |
| | | 0.743 | 0.617 | 0.452 | | (16) |
| GM | 0.503 | 0.574 | 0.617 | 0.452 | 0.404 | |
| ***Albumin (μM)*** | | | | | | |
| | | 602 | 602 | 556 | 511 | (18) |
| | 541 | | 541 | | | (11) |
| | 707 | | | | 526 | (15) |
| | | 561 | 405 | 365 | | (16) |
| GM | 618 | 581 | 510 | 451 | 518 | |

Ref Pop: reference population; CPC: Child-Pugh Class; GM: geometric mean

### 2. Supplemental Results

#### *2.1. Cortisol and cortisone plasma concentrations in individual study participants*

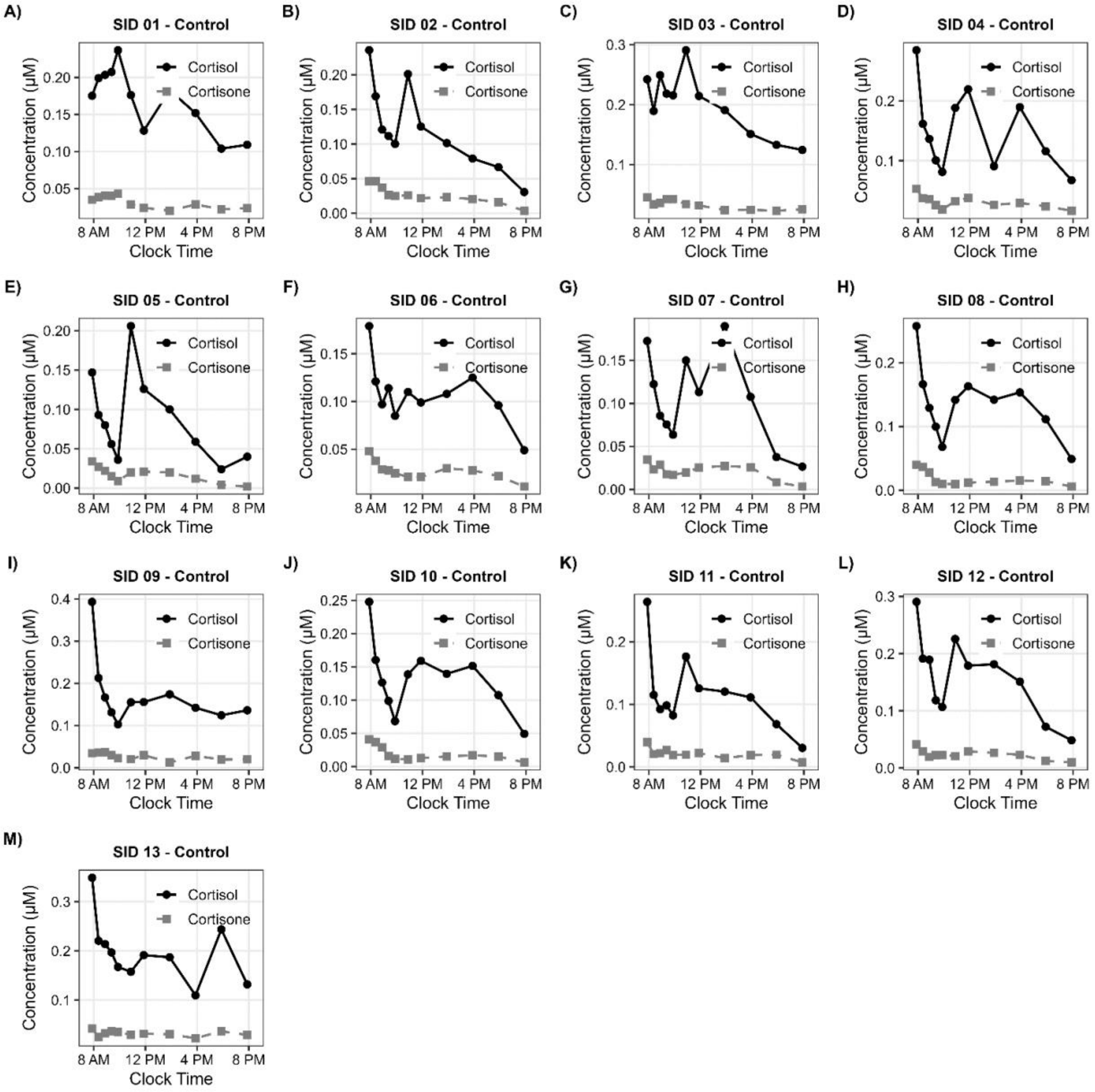

Supplemental Figure 1. Cortisol and cortisone plasma concentration-time curves on control study day for each individual participant (SID).

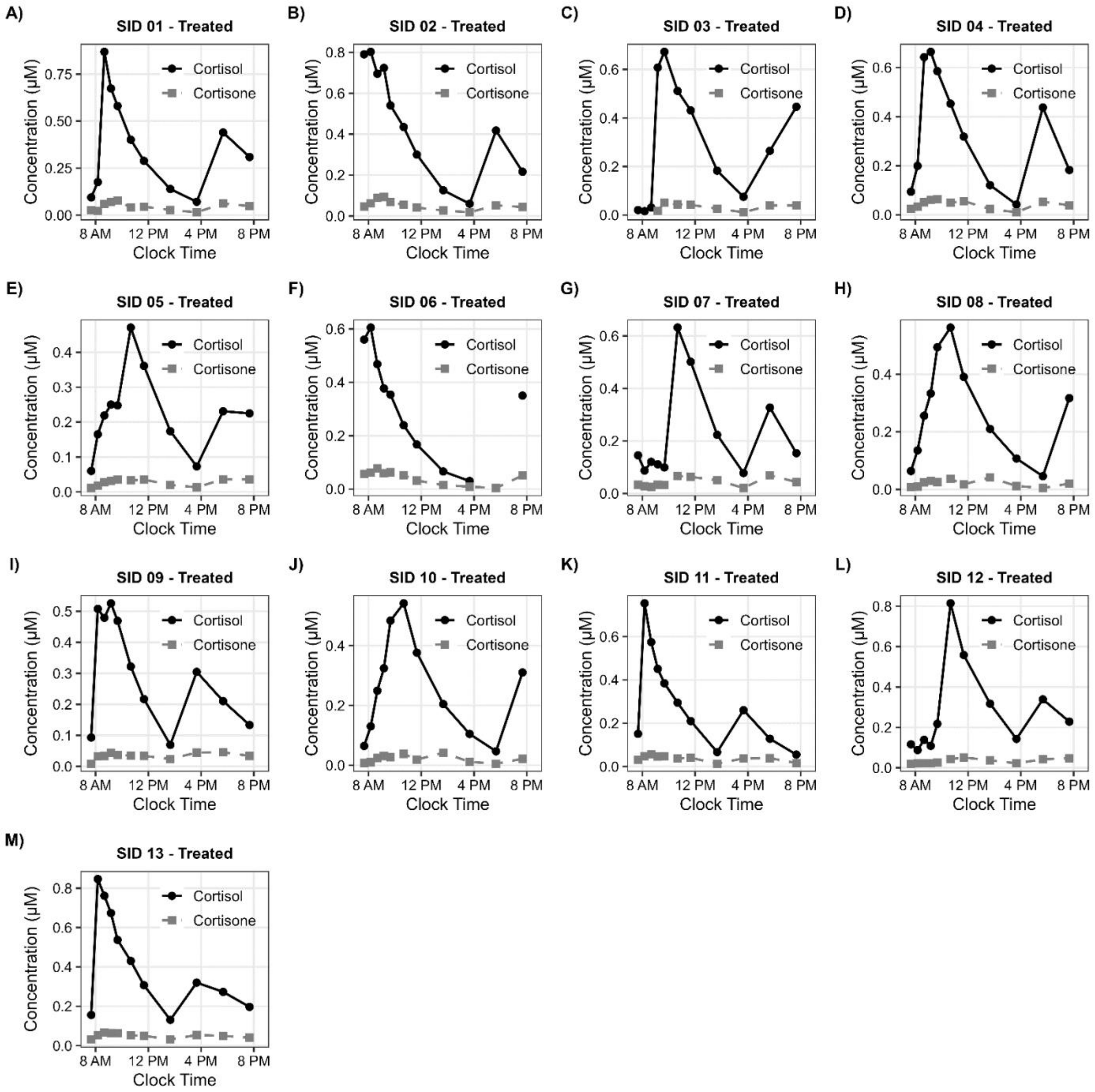

Supplemental Figure 2. Cortisol and cortisone plasma concentration-time curves on treated study day following 7 days synthetic cortisol treatment for each individual participant (SID).

##### *2.2. PPBM 2 best fit disassociation constant values*

Supplemental Table 7. Parameter estimates obtained from the 10x10 bootstrap analysis of the plasma protein binding model (PPBM) 2. Each split represents a bootstrap dataset generated by resampling participants with replacement from the training cohort. Reported values include fit root mean square error (RMSE) and estimated dissociation constants ($K_d$).

| Split | Fit RMSE | $K_{d,\ Cortisol:CBG}$ (μM) | $K_{d,\ Cortisone:CBG}$ (μM) | $K_{d,\ Cortisol:Albumin}$ (μM) | $K_{d,\ Cortisone:Albumin}$ (μM) |
|---|---|---|---|---|---|
| 1 | 2.9 | 0.0122 | 0.146 | 183 | 515 |
| 2 | 3.1 | 0.0109 | 0.165 | 182 | 506 |
| 3 | 3.1 | 0.0120 | 0.173 | 180 | 504 |
| 4 | 2.7 | 0.0116 | 0.151 | 167 | 496 |
| 5 | 2.9 | 0.0103 | 0.150 | 168 | 488 |
| 6 | 2.9 | 0.0121 | 0.153 | 153 | 478 |
| 7 | 3.0 | 0.0119 | 0.160 | 170 | 483 |
| 8 | 2.9 | 0.0124 | 0.157 | 169 | 501 |
| 9 | 2.9 | 0.0178 | 0.158 | 163 | 613 |
| 10 | 3.0 | 0.0146 | 0.138 | 160 | 635 |
| Avg (95% CI) | | 0.0126 (0.0077-0.0174) | 0.155 (0.133-0.178) | 169 (147-191) | 522 (397-647) |

Avg: Average; 95% CI: 95% confidence interval as the product of t-critical value (2.262) and standard deviation

12. Beuschlein F, Else T, Bancos I, Hahner S, Hamidi O, Van Hulsteijn L, et al. European Society of Endocrinology and Endocrine Society Joint Clinical Guideline: Diagnosis and

Therapy of Glucocorticoid-induced Adrenal Insufficiency. The Journal of Clinical Endocrinology & Metabolism. 2024 Jun 17;109(7):1657–83. doi:10.1210/clinem/dgae250
